# Molecular basis for the specific and multivariate recognitions of RNA substrates by human hnRNPA2/B1

**DOI:** 10.1101/144345

**Authors:** Baixing Wu, Shichen Su, Deepak P. Patil, Hehua Liu, Jianhua Gan, Samie R. Jaffrey, Jinbiao Ma

## Abstract

Human hnRNPA2/B1 is an RNA-binding protein that plays important roles in a variety of biological processes, from mRNA maturation, trafficking and translation to regulation of gene expression mediated by long non-coding RNAs and microRNAs. hnRNPA2/B1 contains two RNA recognition motifs (RRM) that provide sequence-specific recognition of widespread RNA substrates including recently reported m^6^A-containing motifs. Here we determined the first crystal structures of tandem RRM domains of hnRNPA2/B1 in complex with various RNA substrates. Our structures reveal that hnRNPA2/B1 can bind two RNA elements in an antiparallel fashion with a sequence preference for AGG and UAG by RRM1 and RRM2, respectively, suggesting an RNA matchmaker mechanism during the hnRNPA2/B1 function. However, our combined studies did not observe specific binding of m^6^A by either the RRM domains or the full-length hnRNPA2/B1, implying that the “reader” function of hnRNPA2/B1 may adopt an unknown mechanism that remains to be characterized.

## INTRODUCTION

Heterogeneous nuclear ribonucleoproteins (hnRNPs) play a variety of roles in regulating transcriptional and post-transcriptional gene expression including RNA splicing, polyadenylation, capping, modification, export, localization, translation, and turnover (Glisovic et al., 2008; Keene, 2007). Each hnRNP contains at least one RNA-binding domain (RBD), such as RNA recognition motif (RRM), K-Homology (KH) domain or arginine/glycine-rich box (He and Smith, 2009). Sequence-specific associations between hnRNPs and their RNA targets are typically mediated by one or more RBDs, which usually bind short, single-stranded RNA (Cook et al., 2011; Gabut et al., 2008), but some also recognize structured RNAs (Auweter et al., 2006).

As a core component of the hnRNP complex in mammalian cells, hnRNPA2/B1 is an abundant protein and has been implicated in widespread biological processes. Gene *HNRNPA2B1* encodes two protein products A2 and B1 through alternative splicing with B1 protein containing an insertion of 36 nucleotides at its N-terminus (Burd et al., 1989; Kozu et al., 1995). Both proteins have an RNA-binding domain (RBD) composed of tandem RRMs separated by a 15-aa linker, a C-terminal glycine-rich region containing a prionlike domain (PrLD) and a nuclear targeting signal (NTS)(Figure 1A).

**Figure 1.**
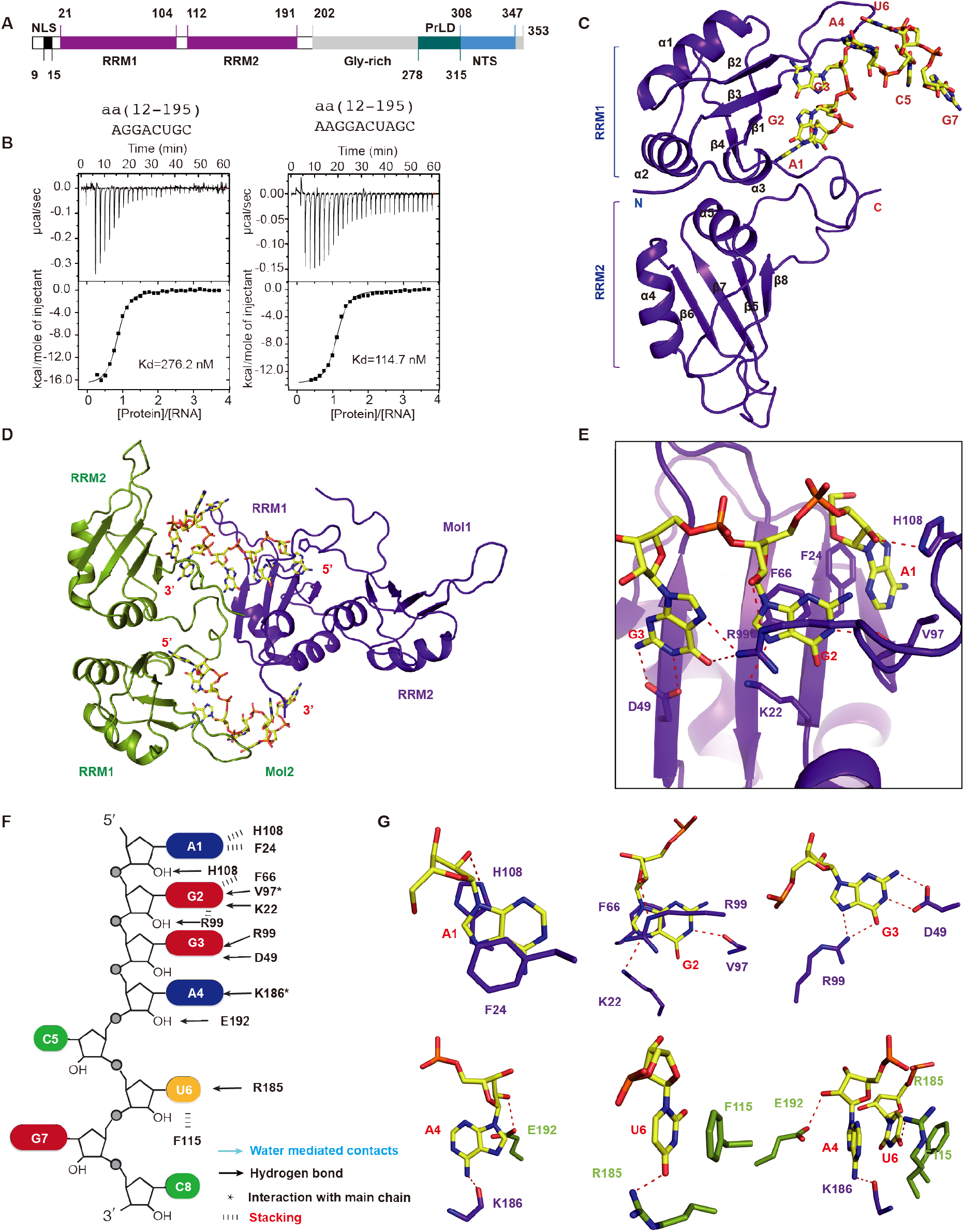
Overview of the structure and ITC of hnRNPA2/B1 in complex with 8mer RNA. (A) Schematic representation of the domain architecture of hnRNPA2/B1. NLS, Nuclear Location Signal; RRM, RNA Recognition Motif; PrLD, Prion-Like Domain; NTS, Nuclear Targeting Signal. (B) ITC results of hnRNPA2/B1(12-195) with 8mer and 10mer RNA targets. Solid lines indicate nonlinear least-squares fit the titration curve, with *ΔH* (binding enthalpy kcal mol^−1^), Ka (association constant) and *N* (number of binding sites per monomer) as variable parameters. Calculation values for K_d_ (dissociation constant) are indicated. [G1](C) Cartoon representation of RRMs in complex with 8mer RNA. The RNA backbone is colored in yellow shown by stick. (D) Molecules from two adjacent asymmetric units. The molecule from another asymmetric unit is colored in green. (E) Intermolecular contacts between RRM1 and 8mer RNA from A1 to G3. (F) Schematic showing RRMs interactions with 8mer RNA 5’-AGGACUGC-3’. (G) Close-up view showing the specific recognition of A1, G2, G3, A4 and U6.[G2]

hnRNPA2/B1 is linked to several biological processes and diseases, especially neurodegenerative disorders. Mutations in PrLD in hnRNPA2/B1 cause multisystem proteinopathy and ALS, through promoting excess incorporation of HNRNPA2/B1 into stress granules and driving the formation of cytoplasmic inclusions in animal models (Kim et al., 2013). hnRNPA2/B1 also regulates hESC self-renewal and pluripotency (Choi et al., 2013).

hnRNPA2/B1 has multiple effects on RNA processing. In addition to participating the whole life cycle of mRNAs, hnRNPA2/B1 is also involved in the activities of many other RNA species. For example, hnRNPA2/B1 can promote association of the long noncoding RNA HOTAIR with the nascent transcripts of HOTAIR target genes, thus to mediate HOTAIR-dependent heterochromatin initiation(Meredith et al., 2016). hnRNPA2/B1 can bind to HIV-1 RNA, causing nuclear retention of the vRNA, as well as microRNA, sorting them into the exosomes.

hnRNPA2/B1 binds consensus motifs in its target RNAs. A transcriptome-wide analysis of hnRNPA2/B1 targets in the nervous system identified a clear preference for UAG(G/A) motifs confirmed by three independent and complementary *in vitro* and *in vivo* approaches (Hutten and Dormann, 2016; Martinez et al., 2016). This is consistent with previous studies indicating that hnRNPA2/B1 binds specifically to UAGGG, GGUAGUAG or AGGAUAGA sequences(Huelga et al., 2012; Ray et al., 2013). Another recent study demonstrated that hnRNPA2/B1 recognizes a consensus motif containing UAASUUAU (S=G or C) in the 3’ UTR of many mRNAs and helps recruiting the CCR4–NOT deadenylase complex (Geissler et al., 2016).

Recently, hnRNPA2/B1 was proposed to bind RNA transcripts containing *N*^6^-methyladenosine, a widespread nucleotide modification in mRNAs and noncoding RNAs (Meyer and Jaffrey, 2014; Meyer et al., 2012). hnRNPA2/B1 was found to mediate m^6^A-dependent nuclear RNA processing events by binding G(m^6^A)C containing nuclear RNAs *in vivo* and *in vitro*. Furthermore, hnRNPA2/B1 was reported to associate with a subset of primary-microRNA transcripts through binding m^6^A, promoting primary-miRNA processing by recruiting the microprocessor complex Drosha/DGCR8 (Alarcon et al., 2015a).

Physiological functions of hnRNPA2/B1 mostly rely on its RNA-binding activity mainly conferred by its RBD. Although a series of studies with different approaches have revealed distinct RNA-binding motifs of hnRNPA2/B1, it is difficult to reconcile these data due to the lack of mechanistic study of RNA selection and binding of hnRNPA2/B1 at the molecular level. Here we report the first crystal structure of the RBD of hnRNPA2/B1 in complex with variant RNA targets. Our results reveal that two RRM subdomains diverge in their RNA-binding properties each having its own preference for an RNA motif, but both having the ability to bind degenerate RNA sequences. In addition, the antiparallel organization of RNA sequences bound to the two RRM domains suggests that hnRNPA2/B1 can promote intra- or inter-molecular association of RNA species, shedding insights into the function of hnRNPA2/B1 in the context of large ribonucleoprotein complexes. Furthermore, our structural data along with detailed RNA-binding measurements did not observe a preference for m^6^A modification by RBD or full-length A2/B1, suggesting that the “reader” function of hnRNPA2/B1 may occur through a yet unknown context *in vivo*, and bioinformatics analysis suggests that hnRNPA2/B1 may bind RNA in response to an “m^6^A switch”, instead of functioning as a direct “reader” of the m^6^A modification.

## RESULTS

### Crystal structure of hnRNPA2/B1 bound to an 8mer RNA substrate

To elucidate the RNA-binding properties of tandem RRMs of hnRNPA2/B1, we purified a number of truncations of the hnRNPA2/B1 protein. Using isothermal titration calorimetry (ITC) method, we characterized the RNA-binding activities of each construct with a set of RNA oligonucleotides of sequences according to previously determined binding motifs. ITC results showed that the construct containing the N-terminal nuclear localization signal (NLS) and two RRM domains, i.e. amino acids (aa) 1-195, can bind target RNAs with high affinity. In addition, deletion of the N-terminal NLS-containing fragment (1-11) had no obvious impact on the RNA binding (data not shown). This suggested that the region containing the tandem RRMs of hnRNPA2/B1 is sufficient for binding target RNA.

We determined the crystal structure of RRMs (aa 12–195) of hnRNPA2/B1 bound to the 8-nt RNA oligonucleotide 5’-A_1_G_2_G_3_A_4_C_5_U_6_G_7_C_8_-3’ (termed 8mer RNA), which is derived from a recent individual-nucleotide-resolution CLIP study (Alarcon et al., 2015a; Martinez et al., 2016). ITC analysis showed that binding of the 8mer RNA occurs at a 1:1 ratio with a *K*_d_ of 276.2 nM (Figure 1B). The crystal structure of RRMs (aa 12-195) in complex with the 8mer RNA molecule was determined to 2.60 Å resolutions, details about data collection and structure refinement are summarized in Table 1. The tandem RRMs and an RNA molecule are in one asymmetric unit (Figure 1C); both RRM domains of hnRNPA2/B1 adopt the characteristic RRM fold, which is a typical β1α1β2β3α2β4 topology consisting of an antiparallel four-stranded sheet adjacent to two helices on the opposite side, similar to previously determined RRM structures of other RNA-binding proteins using both crystallographic and NMR methods (Daubner et al., 2013) (Figure 1C). Each RNA molecule is bound by an RRM1 domain from one hnRNPA2/B1 molecule in an asymmetric unit and an RRM2 domain from another hnRNPA2/B1 molecule in adjacent asymmetric unit (Figure 1D).

### RRM1 specifically recognizes AGG motif

The AGG motif of 8mer RNA substrate is specifically recognized mainly by RRM1 (Figure 1E). For the recognition of the adenine at the first position (A1), the 2’-OH group forms a hydrogen bond and π-π interactions with the side chain of His108. Besides hydrogen bonding interactions, base stacking with the Phe24 on the other side also contributes to the definition of the binding environment (Figure 1F and 1G). The 2’-OH of G2 forms a hydrogen bond with the side chain of Arg99, and N^1^ groups of G2 form hydrogen bonds with the carboxyl group of the main chain of Val97 while N^7^ interacts with the side chain of Lys22. The base of this guanine G2 engages in stacking interactions with the base of the benzene ring of Phe66 and guanidyl group of Arg99 (Figure 1F and 1G). N^1^ and N^2^ of G3, the last nucleotide in the core recognition AGG motif, are hydrogen-bonded to the side chain of Asp49, and O^6^ and N^7^ are recognized by the side chain of Arg99 (Figure 1F and 1G). However, the RNA substrate from A4 to C8 are not well specifically recognized (Figure 1F). The N^6^ of A4 is hydrogen bonded to the main chain of Lys186, and the 2’-OH group is hydrogen-bonded to the side chain of Glu192 (Figure 1F and 1G). The base of U6 is sandwiched between Phe115 and U4 base via π-π stacking, whereas the O^4^ of U6 forms hydrogen bonds with the amino group of the main chain of Arg185 (Figure 1F and 1G).

### Both RRM1 and RRM2 are involved in recognition of the 10mer RNA substrate

The crystal structure of hnRNPA2/B1(12-195) in complex with the 8mer RNA did not provide insight into specific RNA recognition by RRM2. We thus designed another one RNA oligonucleotide shown in Table 2, based on the speculation that RRM2 might recognize UAG according to previous sequencing results (Huelga et al., 2012; Ray et al., 2013). This RNA contains both the AGG motif and the UAG motif. ITC results confirmed that the 10-nt RNA oligo 5’-A_0_A_1_G_2_G_3_A_4_C_5_U_6_A_7_G_8_C_9_-3’ (termed 10mer RNA) has a higher affinity (Figure 1B and Table 2). We successfully obtained the crystal of hnRNPA2/B1(12-195) in complex with the 10mer RNA, which was determined to 1.85 Å resolution (Table 1). Similar to the previous complex structure with 8mer RNA, there is also only one protein molecule and one RNA molecule in the asymmetric unit. However, this time the RNA molecule is recognized by both RRM1 and RRM2 from two hnRNPA2/B1 molecules, of which the other one is from the asymmetric molecule (Figure 2A). The 10mer RNA molecule adopts a single-stranded conformation accommodated into a positively charged groove comprised by the canonical RNA-binding surface of the RRM1 and RRM2 from two hnRNPA2/B1 proteins, in a 5’ to 3’ direction from RRM1 to RRM2 (Figure 2B). Unlike the complex structure of 8mer RNA in which the only RRM1 is involved in specific recognition, the complex structure containing 10mer RNA shows also shows specific recognition by RRM2 (Figure 2C). Interestingly, when RRM1 bound to 8mer and 10mer complex structures are superimposed, the AGG motifs of the two RNA substrates superimpose well. However, the conformation of the rest of the oligos are dramatically different. Notably, the 10mer RNA is more stretched than the 8mer RNA (Figure 2D). Moreover, the root-mean-square deviation (RMSD) of protein backbones between the two structures is merely 0.4 Å, suggesting the protein has not changed substantially when binding the two different RNA targets.

**Figure 2.**
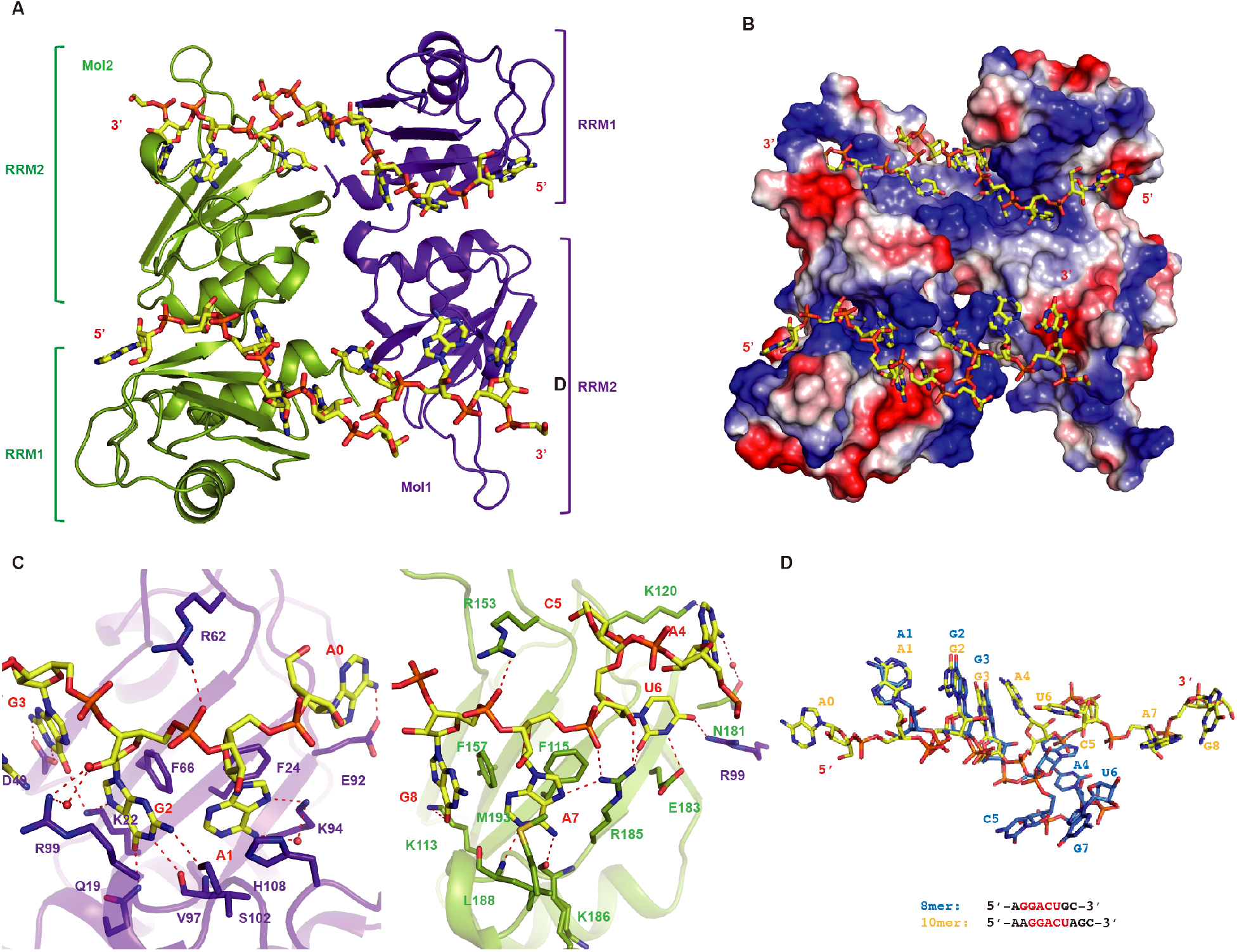
Overview of RRMs in complex with 10mer RNA. (A) Cartoon representation of RRMs in complex with 10mer RNA 5’-AAGGACUAGC-3’. The RNA backbone is colored in yellow shown by stick. The molecule from the adjacent [G1]asymmetric unit is colored in green. (B) Surface representation of RRMs-10mer complex. (C) Intermolecular contacts between RRM1 and residues of 10mer RNA 5’-AAGG-3’, and RRM2 with residues of mer RNA 5’-ACUAGC-3’. (D) Superposition of 8mer and 10mer RNA substrates. The 8mer RNA 5’-AGGACUGC-3’ is colored in marine and the 10mer RNA 5’-AAGGACUAGC-3’ is colored in yellow.

### The RRM2 specifically recognizes UAG motif

Due to the higher resolution, more detailed interactions are observed in the complex structure of 10mer RNA than in the 8mer RNA complex. Though the recognition of the AGG motif in the two structures is quite similar, more specific recognitions of A1 and G2 in the 10mer RNA complex structure are observed. N^1^ of A1 is recognized by the main chain amine group of Val97, whereas N^6^ and N^7^ form hydrogen bonds with the Lys94 side chain either directly or mediated by a water molecule. The base of A1 is clamped by Phe24 and His108 with π-π stacking (Figure 3A and 3B).

**Figure 3.**
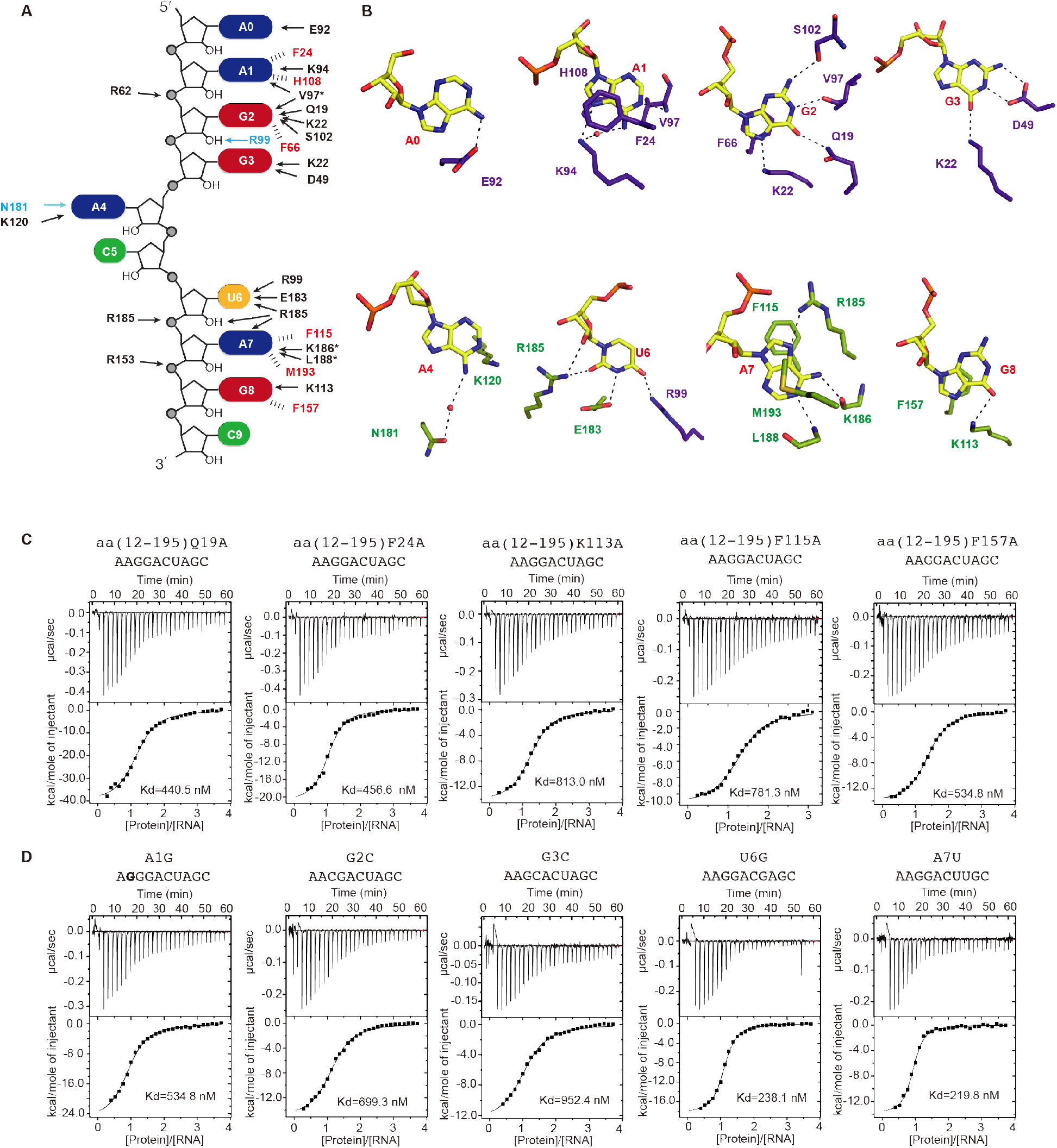
Detailed interactions between RRMs and 10mer RNA. (A) Schematic showing RRMs interactions with 10mer RNA sequence. (B) Close-up view showing the specific recognition from A1 to G8. (C) Mutagenesis study by ITC experiments between protein mutants and 10mer RNA substrate. (D) ITC experiments between hnRNPA2/B1 RRMs and RNA mutants A1G, G2C, G3C, U6G and A7U.

G2 has the most complicated interacting network in this structure, in which the 2’-OH group is hydrogen-bonded to the side chain of Arg99 mediated by a water molecule (Figure 3A). The main chain carbonyl group of Val97 hydrogen bonds to the N^1^, and the side chains of Gln19 and Ser102 cooperatively bond to O^6^ and N^2^ of G2, respectively. N^7^ is hydrogen-bonded to the side chain of Lys22 and exhibits base stacking with Phe66; the phosphate group is hydrogen-bonded to the side chain of Arg62 (Figure 3A and 3B).

N^1^ and N^2^ of G3 are both hydrogens bonded to the side chain of Asp49, and O^6^ of G3 is recognized by the side chain of Lys22 (Figure 3A and 3B). In addition to specific recognition of the AGG motif, N^6^ of the 5’-end extended adenine (A0) is hydrogen bonded to the Glu92, the N^1^, and N^6^ of A4 form hydrogen bonds with the side chain of Lys120 and Asn181 mediated by water, respectively (Figure 3A and 3B).

Unlike the 8mer RNA complex structure, the UAG motif in the 10mer RNA is specifically recognized by RRM2. The side chain of Arg185 hydrogen bonds to both 2’-OH and O^2^ group of U6, and the N^3^ and O^4^ groups are hydrogens bonded to the side chain of Glu183 and Arg99, respectively (Figure 3A and 3B). The base of A7 is also clamped by Phe115 and Met193 through hydrophobic interactions; N^7^ of A7 forms hydrogen bonds with Arg185; N^1^, N^6^, and the phosphate group are hydrogen bonded to the main chain of Leu188, Lys186, and the side chain of Arg185, respectively (Figure 3A and 3B). In addition to the hydrogen bond formed between phosphate group of G8 and the side chain of Arg153, the O^6^ of G8 base is recognized by the side chain of Lys113, and the base of G8 has another π-π stacking with Phe157 (Figure 3A, 3B). The complete structure of C5 and C9 cannot be seen in our structure.

The chains of RRM1 (12-110) and RRM2 (111–195) can be superimposed with a root-mean-square deviation of 0.901 Å with a high sequence identity (Figure S1A). A superimposition of RRM1-AAGG with RRM2-ACUAGC indicated that the recognitions of the AG core motif are very similar in RRM1 and RRM2 (Figure S1B, Figure S1C). We thereafter mutated residues involved in specific recognition of AGG by RRM1 and UAG by RRM2, both of which reduced binding affinities according to ITC (Figure 3C, Figure S2). Although the results of these amino acids mutations lined with expectations, the nucleotide mutations of 10mer RNA, especially the UAG nucleotides recognized by RRM2, have only moderate effects on the binding affinities (Figure 3D, Figure S3, and Table 1).

### Multivariant RNA-binding modes of RRM1 and RRM2

In order to understand the molecular basis for hnRNPA2/B1 recognizing different RNA sequences, we grew the crystals of RBD in complex with different 10mer RNA mutants. Three complex structures containing RNA mutants A1G, U6G and A7U were determined at high resolution (Figure 4A-4C, Table 2). For the A1G mutant (5’-A_0_<u>G</u>_<u>1</u>_G_2_G_3_A_4_C_5_U_6_A_7_G_8_C_9_-3’), the AGG motif is shifted to 5’-end and recognized by RRM1 in a manner almost identical to the wild-type 10mer RNA structure (Figure 4D). In addition, the G3 is recognized by the side chains of Arg99 and Glu18 of RRM1 through hydrogen bond formation (Figure 4D-4F). Therefore, RRM1 of hnRNPA2/B1 can specifically recognize an AGGG motif as demonstrated in the A1G mutant structure. Additionally, RRM2 recognizes UAG in the same manner as was seen in the structure of wild-type 10mer RNA (Figure 4D-4E).

**Figure 4.**
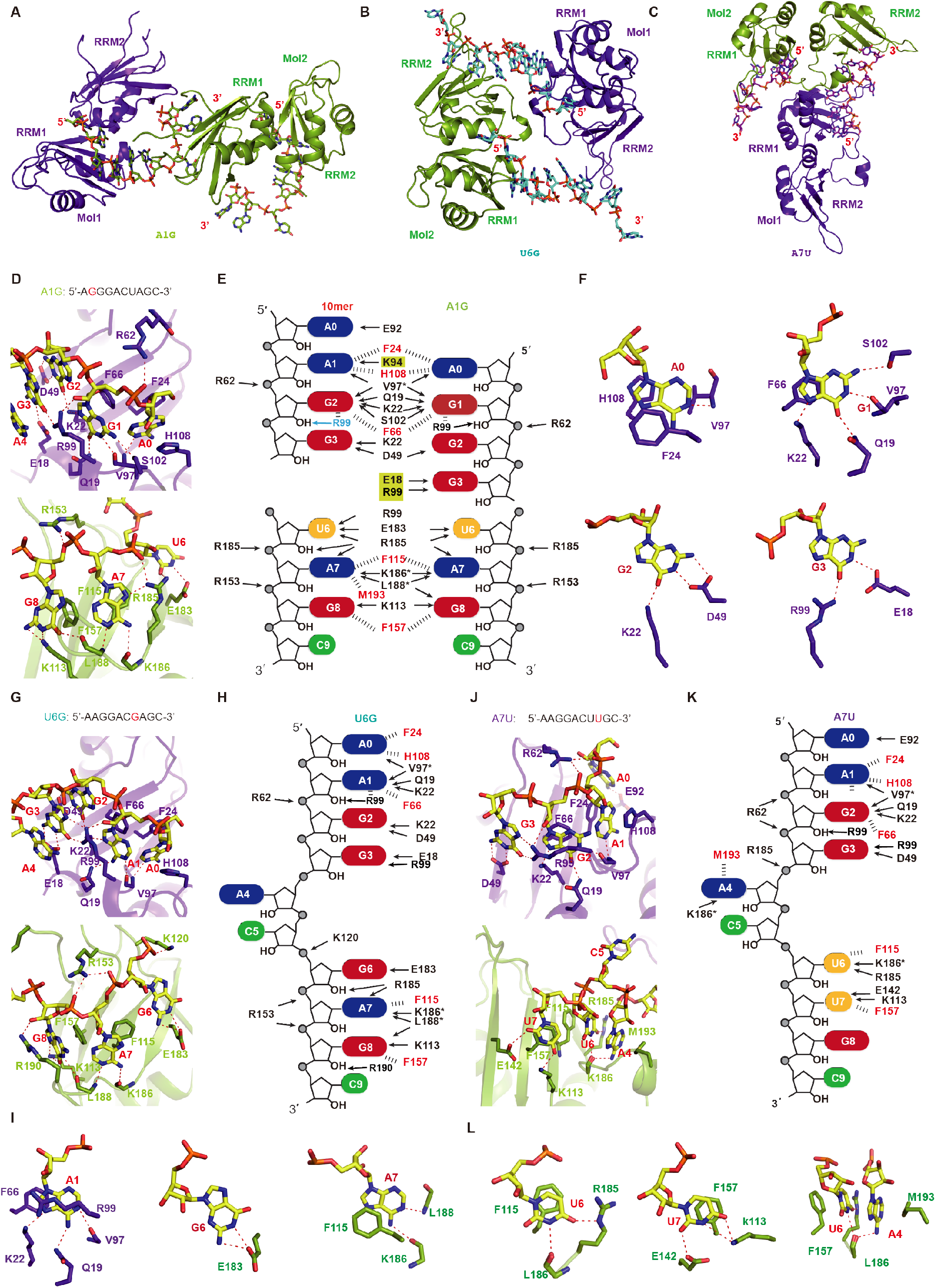
RNA mutants indicate mutivariant binding mode of hnRNPA2/B1 RRMs. (A) Structure of RRMs(12-195) in complex with A1G-RNA 5’-AGGGACUAGC-3’ (B) Structure of RRMs(12-195) in complex with U6G-RNA 5’-AAGGACGAGC-3’ (C) Structure of RRMs (12-195) in complex with A7U-RNA 5’-AAGGACUUGC. (D) Intermolecular contacts between RNA and RRMs in A1G complex. RRM1 is colored in purple-blue, RRM2 is colored in green. (E) Schematic representation of the comparison of different intermolecular interactions between 10mer-RNA 5’-AAGGACUAGC-3’ and A1G-RNA 5’-AGGGACUAGC-3’. (F) Close-up view showing the specific recognition from A1 to G3. (G) Intermolecular contacts between RNA and RRMs in U6G complex. (H) Schematic representation of intermolecular interactions in the U6G complex.[G1] (I) Close-up view showing the specific recognition of A1, G6 and A7 in U6G complex. (J) Intermolecular contacts between RNA and RRMs in A7U complex. (K) Schematic representation of intermolecular interactions in A7U complex. (L) Close-up view showing the specific recognition of U6, U7, and the stacking interactions involved in A4, U6, F157 and M193 in A7U complex.

In contrast, A2/B1 adopts distinct strategies for binding another two RNAs that contained mutations in the UAG motif recognized by RRM2. When U6 is substituted with G in the U6G complex (5’-A_0_A_1_G_2_G_3_A_4_C_5_<u>G</u>_<u>6</u>_A_7_G_8_C_9_-3’ *please check), G6 lost two hydrogen bonds from Arg99 and Arg185, only keeping hydrogen bonds with Glu182. However, the recognition of the AG core motif by RRM2 is still well maintained (Figure 4G-4I). Interestingly, the recognition of AAGG by RRM1 is exactly same as AGGG in the A1G RNA mutant, though their binding modes of AG motif are different, suggesting that RRM1 can accommodate various purine-rich sequences (Figure 4G-4I, Figure S4A, and S4B,). For A7U RNA mutant (5’-A_0_A_1_G_2_G_3_A_4_C_5_U_6_<u>U</u>_<u>7</u>_G_8_C_9_-3’), U7 forms hydrogen bonds with the side chain of Lys113 and Glu142 and forms a π- π stacking interaction with Phe157. More interestingly, U6 adopts a sandwich-like interaction mode with Phe115 and the A4 base, which is exactly same as the U6 in the 8mer RNA substrate (Figure 5J-5L, Figure S4C, and S4D). This suggests that RRM2 can accommodate the pyrimidine-rich UU sequence. Meanwhile, the AAGG recognition by RRM1 in A7U RNA mutant is very similar to 8mer and 10mer RNAs, but different from A1G and U6G RNA mutant.

**Figure 5.**
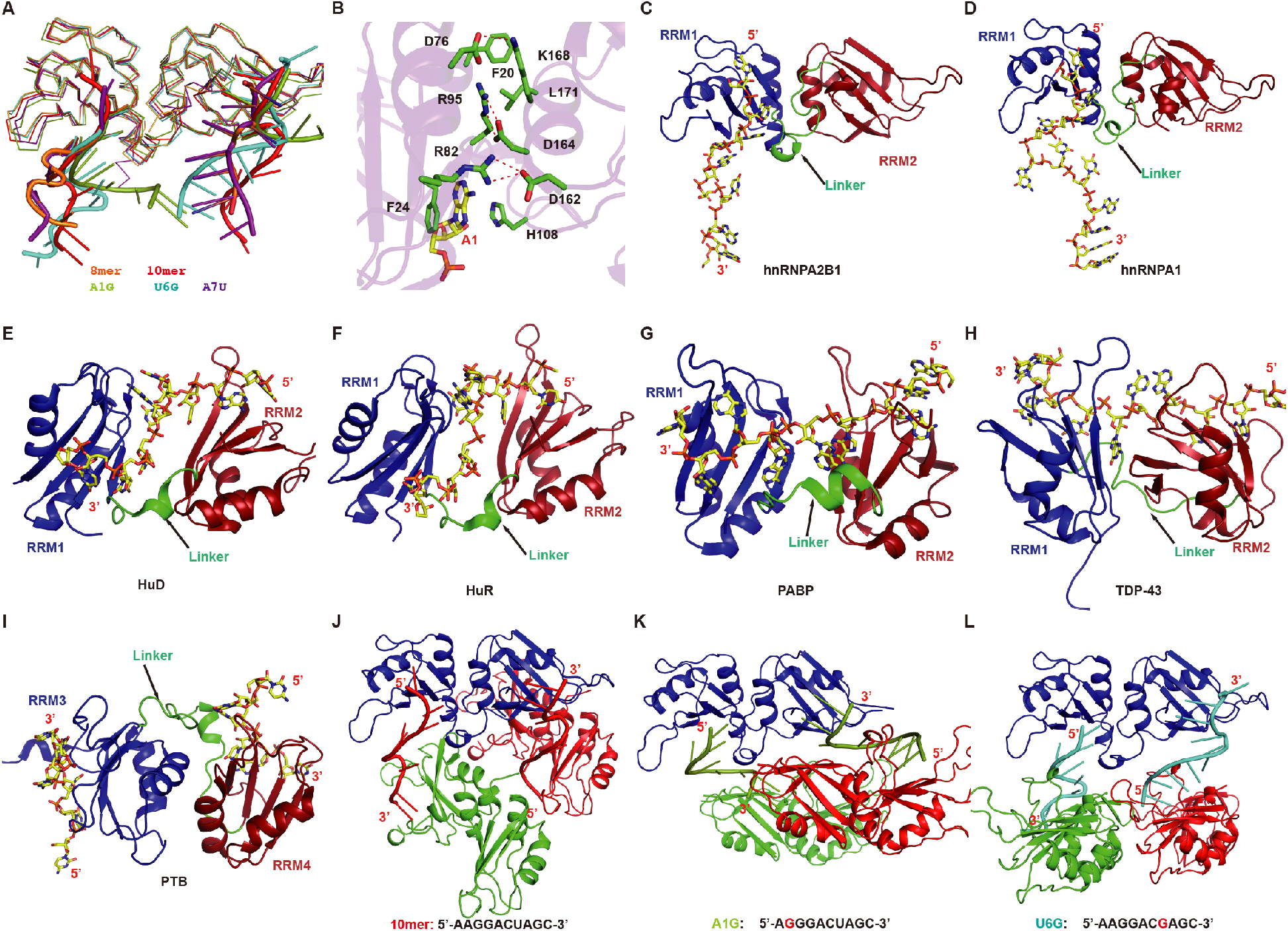
The RNA binding by RRMs adopts an antiparallel mode (a) Superimposition of different structure complex. 8nt-RNA 5’-AGGACUGC-3’ is colored in orange, 10nt-RNA 5’-AAGGACUAGC-3’ is colored in red, A1G-RNA 5’-AGGGACUAGC-3’ is colored in green, U6G-RNA 5’-AAGGACGAGC-3’ and A7U-RNA 5’-AAGGACUUGC are colored in cyan and purple, respectively. (B) Interactions between RRM1 and RRM2, the amino acids participating in interactions are colored in green and RNA is colored in yellow. (C) The overall str[G1]ucture of hnRNPA2/B1-10mer RNA complex. (D) hnRNPA1 in complex with DNA. (E-H) The overall structure[G2] of HuD-RNA, HuR-RNA, PABP-RNA, tdp-43-RNA. From (C) to (H), RRM1 is colored in blue and RRM2 is colored in red, the linker is colored in green pointed out with a black arrow, the RNA backbone is colored in yellow shown by stick. (I) The overall structure of PTB-RNA complex with an antiparallel RNA-binding mode.[G3][G4] RRM3 is colored in blue and RRM2 is colored in red, other labels are the same as from C to H. (J-L) Crystal packing interactions in 10mer, A1G and U6G. To illustrate the detailed packing interactions of hnRNPA2/B1 carrying two antiparallel RNA stands with other hnRNPA2/B1 molecules, three hnRNPA2/B1 molecules and two RNA strands of each complex are selected to show.[G5]

Unlike the effect of mutating the UAG motif recognized by RRM2, which just slightly reduced binding affinities, mutation of the AGG motif recognized by RRM1, such as G2C and G3C RNA mutants, have more obvious effects (Figure 3D). Although we did not obtain crystal structures of these two mutants, our biochemical and structural studies suggested that RRM1 has more stringent recognition for purine-rich AG motif containing RNA sequences, but RRM2 seems to have more broad compatibility to recognize different RNA sequences, including canonical UAG motif, purine-rich GAG and pyrimidine-rich UU sequences.

### hnRNPA2/B1 binds two antiparallel RNA strands

A superimposition of all five structures obtained in this study suggested that hnRNPA2/B1 binds two antiparallel RNA strands using RRM1 and RRM2 concurrently (Figure 5A). The two RRM domains in hnRNPA2/B1, similar to hnRNPA1 (Ding et al., 1999), adopt a synform, in which the linkers between the two domains block the binding of RNA targets by one molecule (Figure 5C and 5D). In contrast, in most known structures of tandem RRM proteins in complex with RNA substrates, the RNAs was bound by only one molecule and the orientation goes from RRM2 to RRM1, such as HuD, HuR, PABP, U2AF65, and TDP-43 (Deo et al., 1999; Lukavsky et al., 2013; Mackereth et al., 2011; Wang et al., 2013; Wang and Tanaka Hall, 2001), and their two RRM domains adopt an anti-form (Figure 5E-5H).

It is notable that there are extensive interactions between RRM1 and RRM2 from the same hnRNPA2/B1 molecule that may account for the synform rigid conformation, including the salt-bridge of Asp76-Lys168, Arg95-Asp164, Arg82-Asp162, and hydrophobic interactions between Phe20 with Leu171 (Figure 5B). The antiparallel orientation of the RNA strands bound by two RRM domains has also been found in the polypyrimidine tract-specific splicing regulator PTB complex structure (Figure 5I), in which the two RRM domains interact each other extensively (Oberstrass et al., 2005). In PTB-RNA complex, the RRM3 and RRM4 form a heterodimer-mediated by a hydrophobic interface, but their RNA binding surfaces point away from each other, with the bound RNAs aligned in an antiparallel orientation, therefore, the PTB RRM3/4 has the capacity to bring together two remote RNA pyrimidine tracts (Oberstrass et al., 2005). In contrast, the binding surfaces of RRM1 and RRM2 in hnRNPA2/B1 do not point away from each other but seem aligned in nearly the same plane. Moreover, the A2/B1 proteins bound to two antiparallel RNA strands can adopt various orientations, as seen in the structures of the 10mer RNA and the two 10mer RNA mutants A1G and U6G (Figure 5J-L, Supplementary Figure 6), mainly due to no direct interactions between the RRM domains bound to the same RNA strands.

### A2/B1 does not specifically recognize m^6^A-modified RNA

In order to assess the hypothesis that A2/B1 may be a direct m^6^A reader as proposed in the previous study (Alarcon et al., 2015a), the m^6^A motif GGACU is included in the 8mer and 10mer RNAs. As shown in the crystal structures, there is no obvious aromatic cage-like surface that can potentially bind the m^6^A nucleotide (Figure 6A-6B), which was shown to be the key m^6^A-specificity element in previous structural studies of YTHDF1, YTHDC1 and MRB1 complexed with GGm^6^ACU (Figure 6C, Figure S6). The crystal structures of hnRNPA2/B1 in complex with GGm^6^ACU could not be obtained. However, we were able to detect binding of hnRNPA2/B1 with the 8mer and 10mer RNA in which an A was replaced with an m^6^A. In both cases, the m^6^A is present within its preferred GGACU sequence context. Notably, the ITC results indicated that the binding affinities of the m^6^A-containing 8mer RNA and 10mer RNA to the tandem RRM (12-195) were reduced one fold and tenfold, respectively, compared to the non-methylated RNA (Figure 6D).

**Figure 6.**
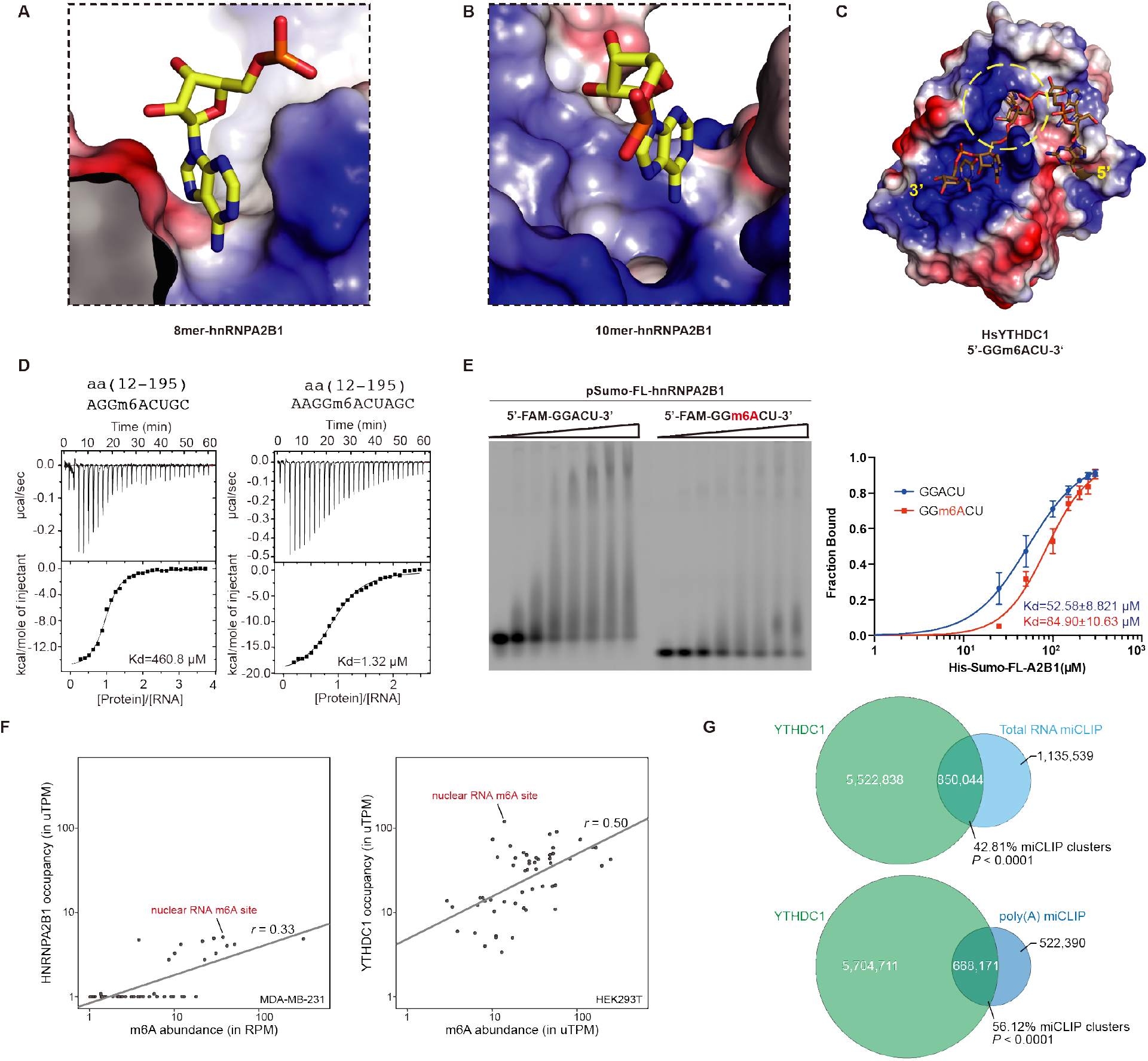
hnRNPA2/B1 does not specifically recognize m^6^A modified RNA. (A) Surface representation of the environment around A4 in 8mer RNA complex. (B) Surface representation of the environment around A4 in 10mer RNA complex. (C) Surface representation of the canonical N^6^-methylated adenosine binding mode in *Hs*YTHDC1. (D) ITC data of hnRNPA2/B1(12-195) with 8mer and 10mer RNA targets carried N^6^-methylated adenosine. (E) EMSA experiment shows the binding affinity of full-length [G1] hnRNPA2/B1 with 5’-FAM-labeled RNA substrates with or without m^6^A modification. (F) YTHDC1 shows preferential binding to m^6^A sites in nuclear RNA compared to hnRNPA2/B1. For hnRNPA2/B1, the m^6^A-Seq reads that overlapped with each m^6^A site was plotted on the x-axis, and the HITS-CLIP reads that overlap with each site were plotted on the y-axis. A similar analysis was used to examine YTHDC1 binding at these m^6^A sites. miCLIP reads that overlapped with the m^6^A sites were plotted on the x-axis, and the YTHDC1 iCLIP reads that overlapped with the m^6^A sites were plotted on the y-axis. (G) YTHDC1-m^6^A tag cluster overlap. A Venn diagram indicating the cluster overlap is shown. Roughly, 43% and 56% of miCLIP tag clusters from total cellular RNA and poly(A) RNA showed a significant overlap with the YTHDC1 iCLIP clusters, respectively.

The N^6^ atoms of A4 in both complex structures of 8mer and 10mer RNAs form hydrogen bonds with A2/B1 directly or through a water molecule, which may provide the possible reason why m^6^A modification would reduce the binding affinities.

However, it could not be excluded that full-length hnRNPA2/B1 may form m^6^A binding cage through its C-terminal fragment, which is not included in our structural study. Therefore, we purified the full-length hnRNPA2/B1 and used it to measure the binding affinities with RNA substrates including the well-known m^6^A motif GGm^6^ACU with 5’-FAM-labeled and a corresponding RNA control lacking the m^6^A modification. We determined the binding affinity by EMSA (Figure 6E, Figure S7). The EMSA analysis indicated that full-length hnRNPA2/B1 has a slightly weaker binding affinity to the RNA with m^6^A modification than the one without m^6^A, suggesting that the full-length hnRNPA2/B1 also does not specifically recognize the m^6^A modified RNA substrates. Together, these different types of experiments suggested that hnRNPA2/B1 does not show enhanced binding to these m^6^A-containing RNAs.

### hnRNPA2/B1 shows binding to only a small subset of nuclear m^6^A sites

To understand hnRNPA2/B1 binding and its relationship to m^6^A, we examined the binding *in vivo* properties of hnRNPA2/B1 on a set of 186 nuclear m^6^A sites comprising m^6^A sites in *XIST, NEAT1*, and *MALAT1*, which have been mapped at single-nucleotide resolution (Linder et al., 2015). In this analysis, we quantified binding at each of these 168 mapped m^6^A residues by assigning each m^6^A residue an ‘intensity value’, which was the normalized number of m^6^A-Seq reads that overlapped each m^6^A residue (Linder et al., 2015). The intensity value is influenced by transcript abundance and m^6^A stoichiometry. We next determined the binding of hnRNPA2/B1 at each of these m^6^A sites from the normalized number of mapped hnRNPA2/B1 HITS-CLIP tags at the m^6^A site. For most m^6^A residues, there was no correlation between m^6^A intensity and hnRNPA2/B1 binding, although 12 m^6^A sites showed proximal hnRNPA2/B1 binding, which might be the result of coincidental proximity between m^6^A and a nonmethylated consensus site recognized by hnRNPA2/B1.

As a control, we analyzed YTHDC1, a nuclear YTH domain-containing m^6^A reader (Patil et al., 2016; Xiao et al., 2016). A similar analysis of YTHDC1 showed increasing YTHDC1 binding with increased m^6^A levels for essentially all m^6^A sites (Figure 6F). Thus, unlike A2/B1, YTHDC1 appears to function as a general nuclear m^6^A reader.

Our finding that only a small subset of nuclear m^6^A sites appear to have bound hnRNPA2/B1 is compatible with the previous analysis by Alarcon *et al*. (Alarcon et al., 2015a), who reported that only 17% of their total m^6^A-seq clusters overlap with the hnRNPA2/B1 tag clusters. To determine if YTHDC1 shows greater overlap with m^6^A than A2/B1 does, we performed a similar cluster overlap analysis. Approximately 43% of the miCLIP clusters from total RNA and 56% clusters from miCLIP of poly(A) RNA (Figure 6G) showed an overlap with YTHDC1 clusters (*P* < 0.0001). Thus, YTHDC1 has a considerably higher overlap with m^6^A than hnRNPA2/B1. Therefore, this analysis again supports the idea that YTHDC1 is the predominant nuclear m^6^A reader compared to hnRNPA2/B1.

## DISCUSSION

The RNA-binding domain of hnRNPA2/B1 comprises two RNA-recognition motifs, RRM1 and RRM2, which is followed by a C-terminal glycine-rich region. hnRNPA2/B1 was previously demonstrated through various analyses to bind UUAGGG and UAG RNA motifs (Hutchison et al., 2002; McKay and Cooke, 1992). Recent CLIP-Seq data further showed that hnRNPA2/B1 prefers to bind A/G-rich sequences. (Huelga et al., 2012). However, these studies did not reveal the molecular basis for the recognition of different RNA substrates of hnRNPA2/B1. Here we determined the crystal structures of the RBD of hnRNPA2/B1 in complex with various RNA substrates, revealing the molecular details of target RNA recognition and providing insights into the potential mechanism for hnRNPA2/B1 in various biological functions.

### Specific and multivariant recognitions of RNA substrates by hnRNPA2/B1

The specific recognition of the AGG motif by RRM1 and the UAG motif by RRM2 imply a role of hnRNPA2/B1 in alternative splicing. In addition, specific recognition of the AG core motif by both RRM1 and RRM2 is in good consistency with previous studies showing that hnRNPA2/B1 can bind A2RE sequence (Munro et al., 1999). Our studies also provided an explanation for the previous study showing that sumoylated hnRNPA2/B1 directs the loading of specific EXO-miRNAs into exosomes by binding GAGG, the so-called EXO motif(Villarroya-Beltri et al., 2013). Furthermore, our results provided the structural basis for hnRNPA2/B1 binding the UA-rich UAASUUAU motif in the 3’ UTR of some mRNAs, which was shown to be necessary for loading the CCR4-NOT complex to mRNAs (Geissler et al., 2016). Taken together, our structures showing sequence-specific RNA-binding properties of hnRNPA2/B1 gives support to previous binding site predictions based on analysis of diverse deep sequencing results (Huelga et al., 2012; Ray et al., 2013) (Table 3).

### Promiscuous binding mode of hnRNPA2/B1

hnRNPA2/B1 shares similar antiparallel arrangements of its bound RNA as PTB, thereby achieving optimal binding of their RNA targets with multiple copies or structured duplexes. Our structural studies of hnRNPA2/B1 offers a molecular basis for the hnRNPA2/B1-HOTAIR interaction which requires multiple nucleotide recognition motifs within HOTAIR (Meredith et al., 2016). These interactions allow hnRNPA2/B1 to preferentially bind to HOTAIR and its targets. The antiparallel orientation of the RNA strands bound by RRM1 and RRM2 in the crystal structures of the hnRNPA2/B1-RNA complex has been reported previously for other tandem RRM domain proteins e.g. polypyrimidine tract-specific splicing regulator PTB (Oberstrass et al., 2005) and zinc finger proteins like MBNL1 (Teplova and Patel, 2008). A similar antiparallel topology in PTB for the RNA binding by RRM3 and RRM4 domains is indicative of a chain reversal loop trajectory for its RNA target (Oberstrass et al., 2005). RNA binding by two or more HNRNPA2B1 molecules may offer the possibility of RNA-templated aggregation and the formation of protein-RNA granules *in vivo* (Martinez et al., 2016).

### Auxiliary role of hnRNPA2B1 in ‘m^6^A switch’

It has recently been demonstrated that hnRNPA2/B1 specifically recognizes m^6^A-modified RNAs (Alarcon et al., 2015a). These RNAs share the m^6^A consensus sequence RGm^6^ACH and directly bind to the m^6^A mark with high affinity *in vivo* and *in vitro* (Alarcon et al., 2015a). Prior to this study, the YTH domain was shown to be a “reader” of m^6^A. However, in addition to directly binding specific proteins, m^6^A can affect RNA binding through an indirect mechanism. This has been shown with two proteins, HuR and hnRNPC, both of which containing RRM domain and do not directly bind m^6^A. In the case of hnRNPC, m^6^A facilitates hnRNPC binding to a UUUUU-tract in mRNAs and long non-coding RNAs (lncRNAs) by promoting local unfolding of RNA. This unfolding is due to the weaker base pairing of U with m^6^A compared to A (Roost et al., 2015). m^6^A-induced RNA unfolding and increased singlestranded RNA accessibility is termed an “m^6^A-switch” (Liu et al., 2015). HuR, also known as ELAVL1, has been found to preferentially bind to the 3’-UTR region of mRNAs that lack m^6^A, increasing their stability and promoting ESCs differentiation (Wang et al., 2014). In this case, m^6^A could impede the formation of a structured RNA motif needed for ELAVL1 binding. Our structural study, combined with biochemistry and bioinformatic results suggests that m^6^A switches may function to enhance hnRNPA2/B1 binding to adjacent non-methylated binding sites, thereby allowing it to regulate miRNA processing. Further *in vitro* and *in vivo* investigations will be required to uncover the details of this mechanism.

## MATERIAL AND METHODS

### Preparation of Protein Samples

Plasmids encoding different fragments of hnRNPA2/B1 were PCR amplified from the human cDNA. PCR products were double digested with restriction endonuclease *BamHI* and *XhoI*, then ligated into a modified pET-28a plasmid carrying the Ulp1 cleavage site. Mutations were generated based on the overlap PCR. Recombinant plasmids were confirmed by DNA sequencing and transformed into *Escherichia coli* BL21 (DE3) to produce target proteins with N-terminal hexahistidine-sumo fusions. *E. coli* cells were cultured in LB medium at 37°C with 50 mg/L kanamycin until the OD_600_ reached 0.6-0.8, then the bacteria were induced with 0.2 mM isopropyl-β-d-thiogalactoside (IPTG) at 18°C for 16 h. Bacteria were collected by centrifugation, resuspended in buffer containing 20 mM Tris-HCl pH8.0, 500 mM NaCl, 20 mM imidazole pH 8.0, and lysed by high pressure. Cell extracts were centrifuged at 18,000 rpm for 1h at 4 °C. Supernatants were purified with Ni-NTA (GE), the target protein was washed with lysis buffer and then eluted with a buffer containing 20mM Tris-HCl, pH8.0, 500mM NaCl and 500mM imidazole. Ulp1 protease was added to remove the N-terminal tag and fusion protein of the recombinant protein and dialyzed with lysis buffer 3 hours. The mixture was applied to another Ni-NTA resin to remove the protease and uncleaved proteins. Eluted proteins were concentrated by centrifugal ultrafiltration, loaded onto a pre-equilibrated HiLoad 16/60 Superdex 75-pg column in an Äkta-purifier (GE Healthcare), eluted at a flow rate of 1 ml/min with the same buffer containing 10mM Tris-HCl pH8.0, 100 mM NaCl. Peak fractions were analyzed by SDS-PAGE (15%, w/v) and stained with Coomassie Brilliant Blue R-250. Purified fractions were pooled together and concentrated by centrifugal ultrafiltration. The concentration was determined by *A_280_*. The protein was concentrated to 10 mg/ml for crystallization trials.

### RNA Oligonucleotides

The RNA oligonucleotides with m^6^A modification were ordered from Dharmacon (Thermo Scientific.), and the others were synthesized by the IDT-394 synthesizer in our own lab. The 5’-FAM-labeled RNA chains were ordered from Bioneer Corporation.

### Crystallization and Data Collection

hnRNPA2/B1 RRMs (12–195) in complex with 8nt-RNA 5’-AGGACUGC-3’ was crystallized using the hanging drop vapor diffusion method by mixing 1μl of protein-RNA mixture (molar ratio 1:1.2) and 1μl of reservoir solution at 20 °C. The crystal suitable for X-ray diffraction was grown in reservoir solution consisting of 0.1M Tris pH8.5 and 25% (w/v) polyethylene glycol 3,350 (Hampton Research). hnRNPA2/B1 RRMs (12–195) in complex with 10nt-RNA 5’-AAGGACUAGC-3’ was screened as above. The crystal suitable for X-ray diffraction was grown in reservoir solution consisting of 0.2M Tri-Sodium Citrate and 20% (w/v) polyethylene glycol 3,350 (Hampton Research). A1G, U6G and A7U were crystallized as the methods mentioned above in solution containing 20% PEG3000, 0.1M Sodium Citrate pH5.5; 20% PEG3350, 0.2M Lithium Sulfate, 0.1M Bis-Tris pH6.5; 25% PEG1500, 0.1M MMT pH9.0, respectively. Data collection was performed at 100 K with cryoprotectant solution (reservoir solution supplemented with an additional 20% (v/v) glycerol). Diffraction data were collected at beamline BL18U of the Shanghai Synchrotron Radiation Facility (SSRF).

### Structure determination and refinement

For hnRNPA2/B1 RRMs (12–195)-8nt complex, the diffraction data set was processed and scaled using HKL3000. The phase was determined by molecular replacement using the program Phaser with the structure of UP1 (PDB code: 1U1Q) as the search model (McCoy et al., 2007). Cycles of refinement and model building were carried out using REFMAC5 and COOT until the crystallography *R*_fector_ and free *R*_free_ converged to 20.3% and 25.6%, respectively (Emsley and Cowtan, 2004; Murshudov et al., 2011). Ramachandran analysis showed that 95.1 of the residues were in the most favored region, with 4.9% in the additionally allowed region. For hnRNPA2/B1-10nt complex, the diffraction dataset was processed and scaled using the HKL3000 package. The phase was determined by molecular replacement using the program Phaser with the hnRNPA2/B1 (12-195) model collected before as the search model. Cycles of refinement and model building were carried out using REFMAC5 and COOT until the crystallography *R*-factor and free *R*-factor converged to 19.0% and 24.3%, respectively. Ramachandran analysis showed that similarly to hnRNPA2/B1-8nt, 99% of the residues were in the most favored region, with 1% in the additionally allowed region. The details of data collection and processing are presented in Table 1. All structure figures were prepared with PyMOL (DeLano Scientific).

### ITC Measurements

ITC assays were carried out on a MicroCal iTC200 calorimeter (GE Healthcare) at 25°C. The buffer used for proteins and RNA oligomers was 10mM HEPES pH 8.0, 50mM KCl, 1mM EDTA, 1mM BME. The concentrations of proteins were determined spectrophotometrically. The RNA oligomers were diluted in the buffer to 5–15 μM. The ITC experiments involved 20-30 injections of protein into RNA. The sample cell was loaded with 250 μl of RNA at 5 μM and the syringe with 80 μL of protein at 100 μM; for weak complexes, the measurement was repeated with increased concentrations. Reference measurements were carried out to compensate for the heat of dilution of the proteins. Curve fitting to a single binding site model was performed by the ITC data analysis module of Origin 7.0 (MicroCal) provided by the manufacturer. ΔG° of protein–RNA binding was computed as RTln(1/KD), where R, T and KD are the gas constant, temperature and dissociation constant, respectively.

### Next-generation sequencing data analysis

Nuclear hnRNPA2/B1 HITS-CLIP sequence data was obtained from a previously published study (Alarcon et al., 2015a) (GEO accession number: GSE70061, SRA accession numbers: SRR2071655 and SRR2071656, last update date: Jun 21, 2015). In addition to the raw data, the author uploaded sequence alignment files GSM1716539_A2B1_HITS_CLIP_1.bedgraph.gz and GSM1716539_A2B1_HITS_CLIP_2.bedgraph.gz were also obtained from the GEO database for comparison purposes. Robust crosslinking-induced mutation sites (CIMS) (FDR ≤ 0.001) in hnRNPA2/B1 HITS-CLIP data were called using a method published elsewhere (Moore et al., 2014). UV-induced deletion sites (Zhang and Darnell, 2011) were used as hnRNPA2/B1-binding sites.

Nuclear m^6^A-seq data from MDA-MB-231 cells were obtained from a previously published study (Alarcon et al., 2015b) (GEO accession number: GSE60213, SRA accession numbers: SRR1539129 and SRR1539130, last update date: Nov 15, 2016). Adapter-free, high-quality sequence reads were aligned to the hg19 genome build using bowtie2 according to the source publication (Alarcon et al., 2015b). RPM (reads per million mapped reads) was calculated using bedtools.

For YTHDC1 binding at m^6^A sites in HEK293T cells, miCLIP sequencing data (Linder et al., 2015) (GEO accession number: GSE63753) and YTHDC1 iCLIP data (Patil et al., 2016) (GEO accession number: GSE78030) was obtained from the GEO database. Sequence alignments were carried out according to the respective publications. Images of genome alignments were prepared using IGV genome browser and Adobe Illustrator.

### Comparison of hnRNPA2/B1 and YTHDC1 binding at m^6^A sites

hnRNPA2/B1 or YTHDC1 binding and m^6^A stoichiometry at 10 bp flanking miCLIP sites (Linder et al., 2015) on nuclear RNAs such as *MALAT1, NEAT1*, and *XIST* was compared using an XY-scatter plot in R. Only m^6^A sites conforming a non-BCANN consensus were considered for this analysis. These represent unique sites obtained from merging (mergeBed -s -d 2) of CIMS- and CITS-based m^6^A site calls from ref. (Linder et al., 2015). All the rRNA, tRNA, and mitochondrial genomic miCLIP sites were removed. Tag counting was performed using the bedtools suite. Tag counts (uTPM+1) were compared using scatter plots and Pearson correlation coefficients (r) were determined in R. Cluster overlap analysis was carried out using bedtools intersect tool (intersectBed -s -u).

## Data Deposition

Coordinates have been deposited in the Protein Data Bank with accession codes for 5EN1 for RRMs-8mer-RNA complex and 5HO4 for RRMs-10mer-RNA structures. A1G, U6G and A7U are 5WWE, 5WWF, 5WWG, respectively.

## Supplementary Materials

Supplementary material is available for this article.

## AUTHOR CONTRIBUTIONS

B.W. and S.S. expressed, purified, and grew crystals of the hnRNPA2/B1-RNA complex and performed the biochemical assays. B.W, S.S., and H.L. synthesized the RNA oligonucleotides. B.W., J.M. and J.G. collected X-ray diffraction data and solved the hnRNPA2/B1-RNA complex structure. D.P. carried out the bioinformatic analysis. B.W., J.M., D. P. and S.J. wrote and revised the manuscript. J.M supervised the structural and biochemical study. S.J. supervised the bioinformatic study.

## ACKNOWLEDGEMENT

We thank the staff of the Beamlines BL18U and BL19U at SSRF for assistance with data collection.

